# A de novo *EDA*-variant in a litter of shorthaired standard Dachshunds with X-linked hypohidrotic ectodermal dysplasia

**DOI:** 10.1101/397372

**Authors:** Danae Vasiliadis, Marion Hewicker-Trautwein, Daniela Klotz, Michael Fehr, Stefka Ruseva, Jennifer Arndt, Julia Metzger, Ottmar Distl

**Affiliations:** Institute of Animal Breeding and Genetics, University of Veterinary Medicine Hanover, Foundation, 30559 Hanover, Lower Saxony, Germany; Institute of Pathology, University of Veterinary Medicine Hanover, Foundation, 30559 Hanover, Lower Saxony, Germany; Clinic for Small Animals, University of Veterinary Medicine Hanover, Foundation, 30559 Hanover, Lower Saxony, Germany

**Author notes:** Corresponding author: Ottmar Distl, Institute of Animal Breeding and Genetics, University of Veterinary Medicine Hanover, Bünteweg 17p, 30559 Hannover, Germany.

**Keywords:** ectodermal dysplasia, canine, whole-genome sequencing, X-linked inheritance, Ectodysplasin-A

## Abstract

In this study, we present a detailed phenotype description and genetic elucidation of the first case of X-linked hypohidrotic ectodermal dysplasia in the shorthaired standard Dachshund. This condition is characterized by partial alopecia, missing and malformed teeth and a lack of eccrine sweat glands. Clinical signs including dental X-raying and histopathological findings were consistent with an ectodermal dysplasia. Pedigree analysis supported an X-recessive mode of inheritance. Whole-genome sequencing of one affected puppy and his dam identified a 1-basepair deletion within the *ectodysplasin-A* gene (CM000039.3:g.54509504delT, PRJEB27789). Sanger sequencing of further family members confirmed the PRJEB27789-variant. Validation in all available family members, 37 unrelated shorthaired standard Dachshunds, 128 Dachshunds from all other breeds and samples from 34 dog breeds revealed the PRJEB27789 variant to be private for this family. Two heterozygous females showed very mild alopecia but normal dentition. Since the dam is demonstrably the only heterozygous animal in the ancestry of the affected animals, we assume that the PRJEB27789-variant arose in the germline of the granddam or in an early embryonic stage of the dam. In conclusion, we detected a very recent de-novo *EDA* mutation causing X-linked hypohidrotic ectodermal dysplasia in the shorthaired standard Dachshund.

## Introduction

X-linked hypohidrotic ectodermal dysplasia (XLHED) is a hereditary defect that causes malformations of structures that emerge from the ectoderm, including hair, teeth and sweat glands (Söderholm and Kaitila 1985; Clarke 1987; Kere et al. 1996; Itin 2014). XLHED manifests in humans with alopecia, a lack of sweat glands and possibly other exocrine glands, oligodontia and conically shaped teeth (Clarke et al. 1987; Cluzeau et al. 2011). The affected individuals therefore suffer from hypohidrosis, which can lead to hyperthermia (Mills 1968). They are also affected with decreased mucociliary clearance, predisposing to respiratory tract infections, decreased lacrimal function and dry eye, and problems with mastication (Beahrs et al. 1971; Söderholm and Kaitila 1985; Clarke et al. 1987; Gilgenkrantz et al. 1989; Johnson et al. 2002; Casal et al. 2007). In humans, XLHED is the most frequent form of ectodermal dysplasias (Kere et al. 1996). Ectodermal dysplasias (ED) are a group of genetically heterogeneous diseases. XLHED is caused by alterations in the *ectodysplasin-A (EDA)* gene (Kere et al. 1996), but there are also autosomal dominant, recessive and unknown mutations that cause an ED phenotype (Pinheiro and Freire-Maia 1994; Munoz et al. 1997; Ho et al. 1998; Monreal et al. 1999). In humans, only four genes (*EDA, ectodysplasin-A receptor (EDAR), EDAR-associated death domain (EDARADD)*, and *wingless-type MMTV integration site family, member 10A (WNT10A)*) are causative for 90% of hypo-/anhidrotic ED cases (Cluzeau et al. 2011).

*EDA* is located on the X-chromosome. It has several transcripts and protein isoforms. In humans, the two isoforms EDA-A1 and EDA-A2 interact with EDAR and ectodysplasin-A2 receptor (EDA2R, formerly XEDAR), respectively (Yan et al. 2000). EDAR can activate nuclear factor kappa-B (NF-κB), c-Jun N-terminal kinase (JNK) and cell death pathways (Kumar et al. 2001; Koppinen et al. 2001). The EDAR-activation of the NF-κB pathway is needed for the development of skin appendages (Mikkola and Thesleff 2003; Schmidt-Ullrich et al. 2006; Cui and Schlessinger 2006).

Many ED variants were identified in different species. More than 80 causal mutations are described in humans (Huang et al. 2015). Cases in other species include mice (Srivastava et al. 1997), cattle (Drögemüller et al. 2001), horse (Ramzan et al. 2001), fish (Kondo et al. 2001) and dogs. ED phenotypes were shown for the dog breeds Bichon Frisé (Grieshaber et al. 1986), Labrador Retriever (Kunkle 1984), Miniature Poodle (Selmanowitz et al. 1970b), Whippet (Thomsett 1961), Cocker Spaniel (Kral and Schwartzman 1964), Basset Hound (Chastain and Swayne 1985), mixed breeds of miniature Pinscher and Pekingese (Moura and Cirio 2004), German Shepherd (Casal et al. 1997) and mixed breed dogs (Waluk et al. 2016). ED phenotypes have been suspected in the breeds Belgian Shepherd, Beagle, Rottweiler, Yorkshire Terrier, French Bulldog and Lhasa Apso (Miller et al. 2013). In those latter dog breeds, the authors state that it was not always possible to determine whether the dogs exclusively had alopecia or expressed further ectodermal defects, too. The hairless dog breeds Xoloitzcuintle and Chinese crested dog also exhibit an ED phenotype, as they frequently have missing teeth. The hairlessness in these breeds is caused by an autosomal dominant defect in the *FOXI3* gene (Drögemüller et al. 2008). The clinical signs of XLHED-affected dogs are almost identical to those of affected humans (Casal et al. 2007). Treatment of the disease for mice is possible using prenatal intravenous administration of recombinant EDA (Gaide and Schneider 2003). Dogs, which were treated postnatally, had improved permanent dentition and normalized lacrimation, as well as improved resistance to airway infections and sweating ability (Casal et al. 2007; Mauldin et al. 2009). Recently, human children were treated prenatally with promising results, too (Schneider et al. 2018). The objective of this study was to characterize the ED phenotype in a litter of shorthaired standard Dachshunds, to determine the most likely mode of inheritance after collection of records of relatives, and to unravel the responsible mutation. Whole-genome sequencing data were used to screen the genome for ED-associated variants and then, we validated the ED-associated variant in the close relatives of the affected Dachshunds, a large panel of different Dachshund breeds and other dog breeds.

## Materials and Methods

### Ethics Statement

All animal experiments were performed according to national and international guidelines for animal welfare. All dogs in this study were privately owned and written informed consent of their owners was obtained for this specific study. The Lower Saxony state veterinary office at the Niedersächsisches Landesamt für Verbraucherschutz und Lebensmittelsicherheit, Oldenburg, Germany, was the responsible Institutional Animal Care and Use Committee (IACUC) for this study. This particular study was approved by the Niedersächsisches Landesamt für Verbraucherschutz und Lebensmittelsicherheit (LAVES) under the file number 33.8-42502-05-18A294. Further dog samples of different breeds used as controls were taken from the biobank of the Institute of Animal Breeding and Genetics at the University of Veterinary Medicine Hanover.

### Animals

Four euthanized male shorthaired standard Dachshund puppies (dogs A, B, C and D) with an ED phenotype were presented at the Institute for Animal Breeding and Genetics of the University of Veterinary Medicine Hanover. The owner reported that the affected puppies developed nasal discharge and respiratory distress, which is why the puppies were euthanized at the age of eight days. The other three puppies were female and showed no signs of a disease. The litter was bred according to the Fédération Cynologique Internationale (FCI) rules within the Deutscher Teckelklub 1888 e.V. (DTK). Pedigrees officially issued by the DTK for the litter with the affected puppies, their parents, grandparents and further relatives were provided by the dog owner.

### Diagnostic imaging, necropsy and histo-pathology

The four affected, eight-day-old male shorthaired standard Dachshund puppies were clinically examined. Three of the four affected puppies (dog B, C and D) were X-rayed to evaluate their dentition. One puppy (dog A) was submitted for necropsy and histo-pathological examination. Tissue samples were fixed in 10% neutral buffered formalin before being embedded in paraffin. For histological examination, 2–3 μm thick sections were cut and stained with hematoxylin and eosin. To compare the results with normal control tissue, formalin-fixed tissue was obtained from a 9-day-old puppy that had to be euthanized due to another reason.

### DNA and RNA sampling and extraction

We took tissue samples of the tails from the four affected puppies and EDTA-blood samples from the dam (dog E) and maternal granddam (dog F). In addition we collected EDTA-blood samples from three female full-siblings of the affected puppies (dog H, I, and J) and two full-siblings of the dam (dog K and L). DNA samples from the maternal great-granddam (dog G) and four paternal half-siblings of the dam (dogs M, N, O, P) were already in our biobank. Genomic DNA extraction was performed using a standard salting out procedure using chloroform. Hair root samples from the dam (dog E) were taken and stored in RNAlater for RNA isolation using RNeasy Lipid Tissue Mini Kit (Qiagen, Hilden, Germany). Reverse transcription reactions from the RNA samples of the dam and a Great Dane reference animal were performed with Invitrogen Super Script IV ReverseTranscriptase (Thermo Fisher Scientific, Waltham, MA, USA).

### Whole genome sequencing (WGS)

We prepared libraries with the NEBNext Ultra II DNA Library Prep Kit for Illumina (New England BioLabs, Ipswich, MA, USA). The libraries of one affected puppy and his dam (dog A and E) were sequenced on an Illumina NextSeq500 (Illumina, San Diego, CA, USA) using the 2×150bp in paired-end mode. Quality control was applied with fastqc 0.11.5 (Andrews 2010) and reads were trimmed using PRINSEQ (V 0.20.4) (Schmieder and Edwards 2011). We mapped the reads to the canine reference genome CanFam 3.1 (GCA_000002285.2, Ensembl, www.ensembl.org) using BWA 0.7.13 (Li and Durbin 2009). Sorting, indexing and marking of duplicates was performed using SAMtools 1.3.1 (Li et al. 2009) and Picard tools (http://broadinstitute.github.io/picard/, version 2.9.4). We employed GATK version 4.0 (McKenna et al. 2010) using Base Quality Score Recalibration (BQSR), Haplotype Caller and Variant Recalibrator for variant calling. Variant effect estimation was performed on basis of SNPEff v 4.1 (Cingolani et al. 2012) for predictions on CanFam3.1.86 database. For comparison, WGS data from five dogs of the breeds Briard (n=2), German Drahthaar (n=3) and a Chinese Wolf (SRX1137188) were available. We searched the WGS data for hemizygous mutant variants on the X-chromosome and homozygous mutant variants on all autosomes using SAS, version 9.4 (Statistical Analysis System, Cary, NC, USA). Only those variants, which were predicted to have high to moderate effects using the prediction toolbox SNPEff were chosen for further analysis. Additionally we used the Ensembl Variant Effect Predictor (McLaren et al. 2010), with CanFam 3.1 as reference genome, for SIFT predictions (Kumar et al. 2009). We also applied the PolyPhen-2 (Polymorphism Phenotyping v2) tool for variant effect predictions (Adzhubei et al. 2010). The results were compared with a candidate gene list for ectodermal dysplasia (Table S1). Only variants in those genes exclusively found in dog A were considered for further analysis.

### Validation

In total, 12 shorthaired standard Dachshunds closely related with the affected puppies, further 37 unrelated shorthaired standard Dachshunds and 128 Dachshunds from other varieties (shorthaired, wirehaired and longhaired in the sizes standard, miniature and rabbit) as well as 70 dogs of 34 different breeds from our biobank were used as reference animals. A restriction fragment length polymorphism (RFLP) for mutation detection of the PRJEB27789 variant within *EDA* was developed using the restriction enzyme *Alu*I (New England Biolabs) (Table S2). We designed primers for the polymerase chain reaction (PCR) using Primer3 (http://bioinfo.ut.ee/primer3/) (Untergasser et al. 2012) (Table S2). The reaction volume of 20 µl was composed of 2 µl DNA or cDNA, 11.8 µl H2O, 4.4 µl Q-Solution (Qiagen), 2.2 µl PCR-buffer (Qiagen), 0.5 µl dNTP mix, 0.3 µl (5 u/µl) Taq DNA Polymerase (Qiagen), 0.5 µl forward and 0.5 µl reverse primers. The reaction was performed on a SensoQuest Labcycler thermocycler (SensoQuest, Göttingen, Germany) setting 5 minutes denaturation at 95°C, 34 cycles of 94°C for 45 seconds, 61.3°C annealing temperature, 72°C for 45 seconds, a final extension step at 72° for 10 minutes and then 4°C for 5 minutes. The incubation of PCR amplificates (15 µl) with *Alu*I (0.5 µl) was performed with 2.5 µl H2O and 2 µl Cutsmart buffer (New England Biolabs) at 37°C for 12 hours. Products were separated by gel electrophoresis using 4% agarose gels (Rotigarose, Carl Roth, Karlsruhe, Germany). Genotypes were determined by visual examination under UV illumination (BioDocAnalyze, Biometra, Göttingen, Germany). In addition, we used Sanger sequencing to confirm the *EDA*-mutation (Table S2). PCR and cDNA-PCR products were treated with exonuclease I and FastAP Thermosensitive Alkaline Phosphatase and then sequenced on an ABI 3730xl DNA Analyzer (Eurofins GATC Biotech, Konstanz, Germany). We analyzed the Sanger sequence data with Sequencher 4.8 software (GeneCodes, Ann Arbor, MI, USA). ORFfinder (NCBI, https://www.ncbi.nlm.nih.gov/orffinder/) was used to predict the effects of the mutation on the open reading frame. The wildtype and mutated predicted protein sequences were compared using Clustal Omega (https://www.ebi.ac.uk/Tools/msa/clustalo/) (Sievers et al. 2011). The position of the protein domains of ectodysplasin were assessed with Ensembl and Superfamily (Gough et al. 2001), Gene3D (Lewis et al. 2018), Prosite_profiles (Sigrist et al. 2013) and Pfam (Finn et al. 2016) as domain sources.

### Data availability

The whole-genome sequencing data of the affected puppy and dam were deposited in the NCBI Sequence Read Archive (http://www.ncbi.nlm.nih.gov/sra) under the SRA accession number PRJNA309755 (SAMN09435548 and SAMN09435549). The *EDA*-variant is submitted under the accession number PRJEB27789 in the European Variant Archive (EVA). Supplemental material is available at Figshare: EDA_Vasiliadis_Dachshund_Ectodysplasin_de_novo.

## Results

### Clinical presentation

All four male standard shorthaired standard Dachshund puppies of the litter showed alopecia in the area of the skullcap, the median area from the base of the tail up to the last ribs, the lower chest and abdomen and the upper inner part of the limbs (Figure 1). The puppies had normal weight at birth. The parents of the affected puppies were phenotypically inconspicuous and officially, FCI-registered standard sized shorthaired Dachshund breeding dogs (Figure 2). Two of the female puppies of the litter (dogs I and J) had mild hypotrichosis of the forehead, ears and lower chest. The female puppies showed no other abnormalities and thrived well.

**Figure 1:**
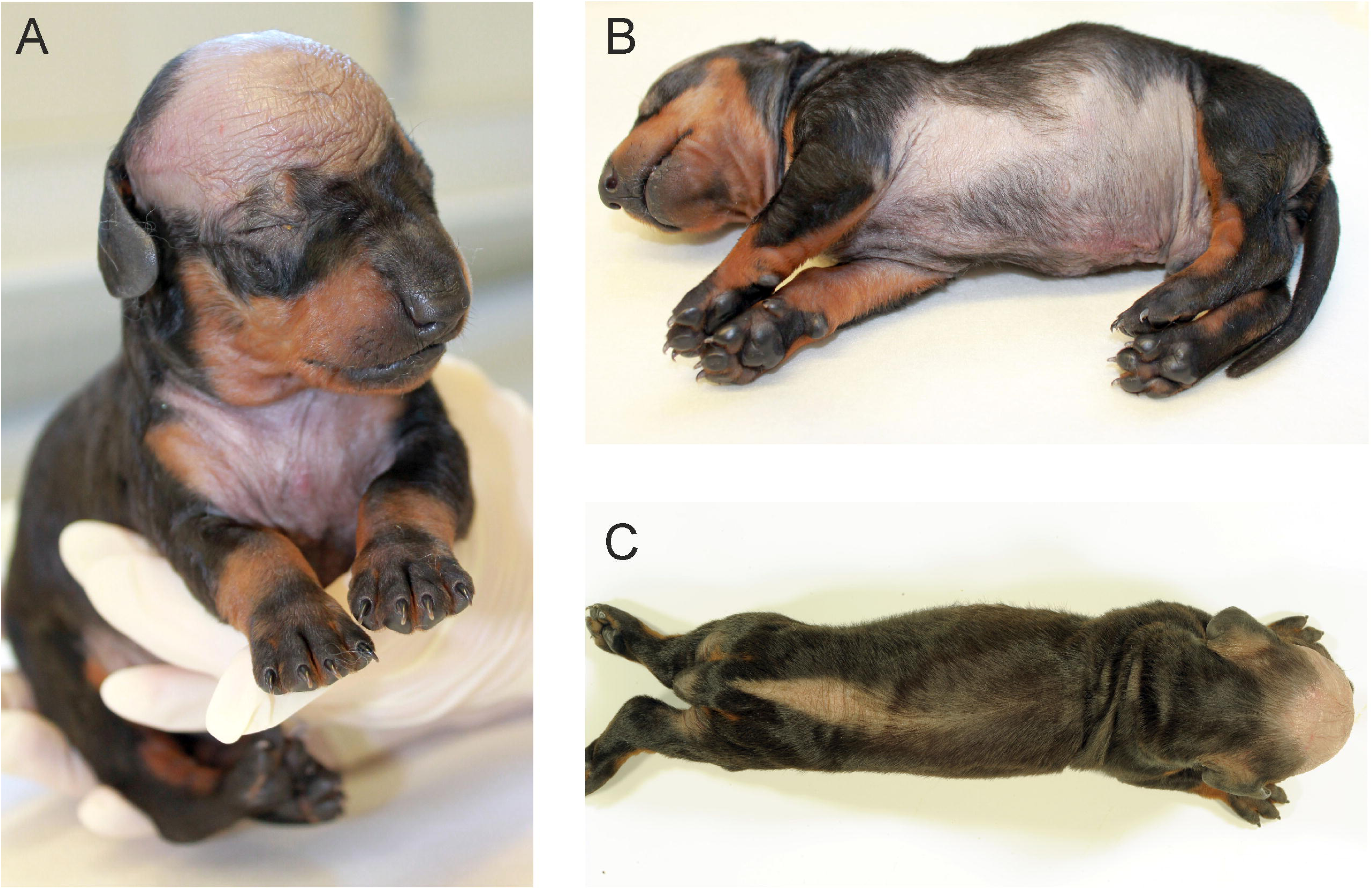
Phenotypic appearance of a male affected shorthaired standard Dachshund (dog A). The characteristic alopecia pattern is easily recognizable. (A) The forehead and the chest with the well demarked hypotrichosis. (B) The whole underbelly is hairless, too. Note that the nails and footpads are developed normally. (C) The alopecia also extends in a dorsal stripe from the base of the tail up to the last ribs. The three other affected male puppies shared the exact phenotype. Three of the puppies were black and tan and one was brown.

**Figure 2:**
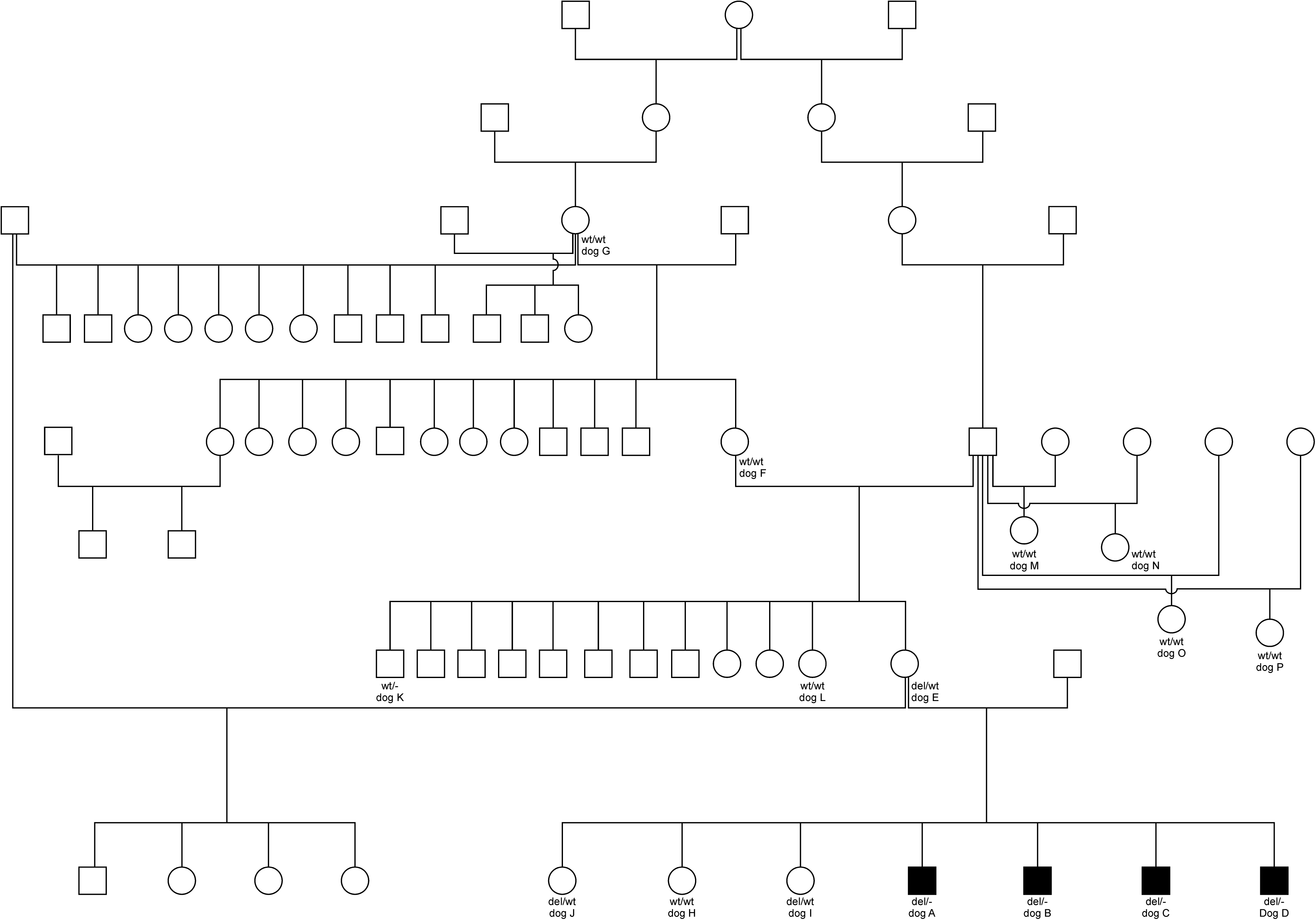
Pedigree of the affected shorthaired standard Dachshund family. Family members, which are mentioned in this study, are marked with the letters A to P. Also given are the genotypes for every dog we were able to get DNA samples from. A square symbol means a male individual, and a circle a female. Filled out symbols mark the XLHED-affected individuals.

### Necropsy and histopathological examination

Necropsy and histo-pathological examination was done in one male puppy (dog A). Aside from the alopecia restricted to specific body areas, there were no macroscopic pathological findings. Histologically, there was a focally reduced number and lack of hair follicles in the alopecic areas of the head and back. A medium degree, focal, irregular hyperplasia of the epithelium was present in the skin of the head. There also was a medium degree, focal, subacute, purulent folliculitis as well as a low-grade inflammatory infiltration of both epidermis and dermis. The skin of the footpads missed sweat glands, and the trachea, esophagus and bronchia were devoid of glands. There were no cilia detectable in the trachea, but in the bronchial epithelium. There was no indication of bronchopneumonia (Figure 3).

**Figure 3:**
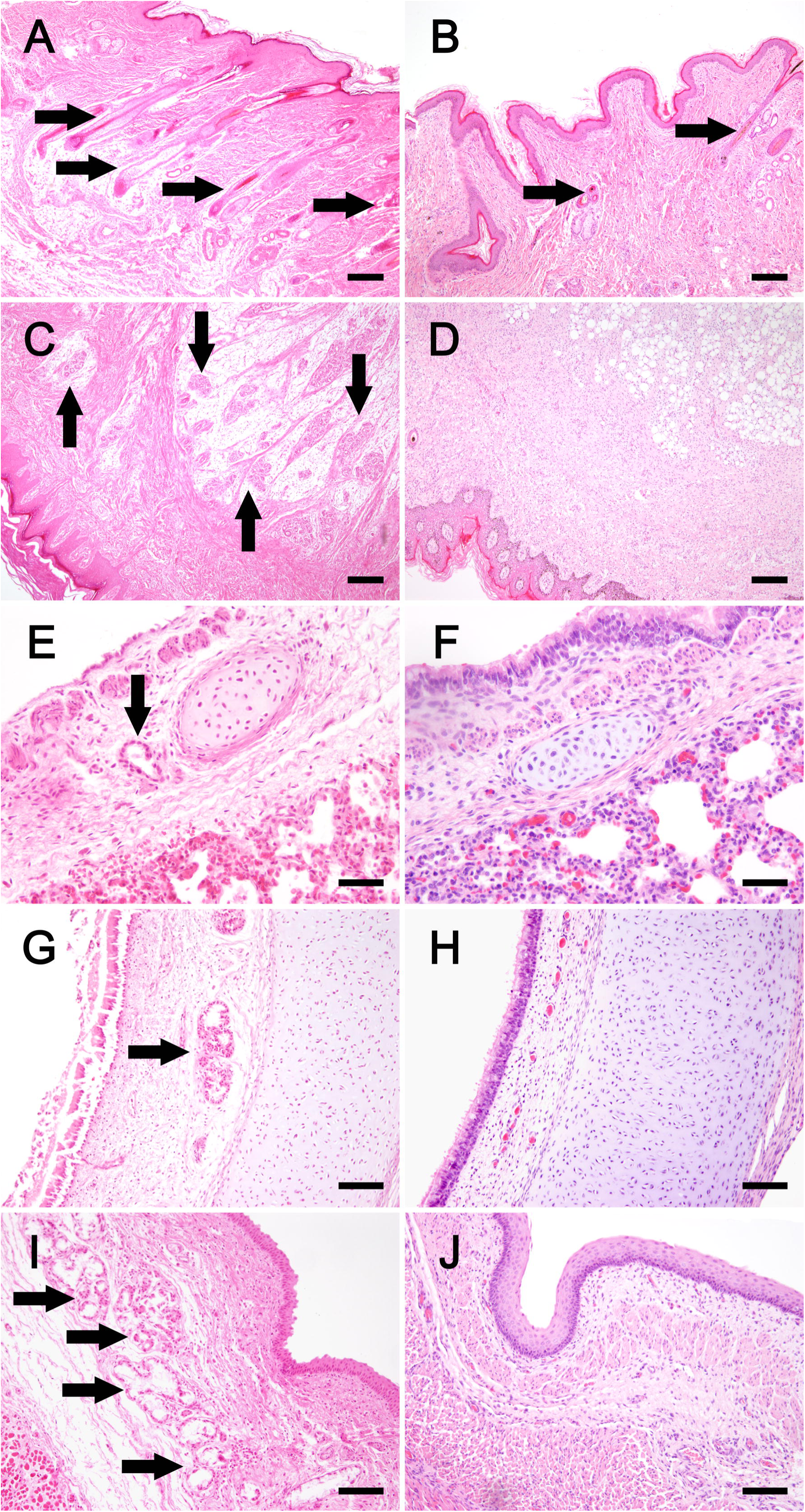
Histopathology of the eight days old Dachshund puppy (right panels) compared to a 9-days-old control puppy (left panels). In normal skin, numerous hair follicles are visible (A), while in alopecic areas the hair follicle number is highly reduced with a segmental absence (B). Numerous eccrine sweat glands in the fat pad can be detected in the control footpad (C), whereas in the examined Dachshund puppy no glandular structures are recognizable (D). Individual bronchial glands can often be observed next to cartilage structures (E), whereas in the examined Dachshund puppy no glandular structures were detected in the bronchi (F). In the submucosa of the trachea (G) and esophagus (I), numerous nests of glands are usually recognizable, while they are completely absent in the examined Dachshund puppy (H, J). (Hematoxylin and eosin stains; Bar in A, B, C, D: 200µm; Bar in E and F: 50 µm; Bar in G, H, I, J: 100 µm).

### Diagnostic Radiology

The full-body radiological images of three affected puppies (dogs B-D) revealed no anomalies of the skeleton. Dental imaging of two of the affected puppies (dog B and D) showed multiple missing premolars and irregularly spaced teeth. However, dog C had a complete dentition and even an additional premolar (Figure 4).

**Figure 4:**
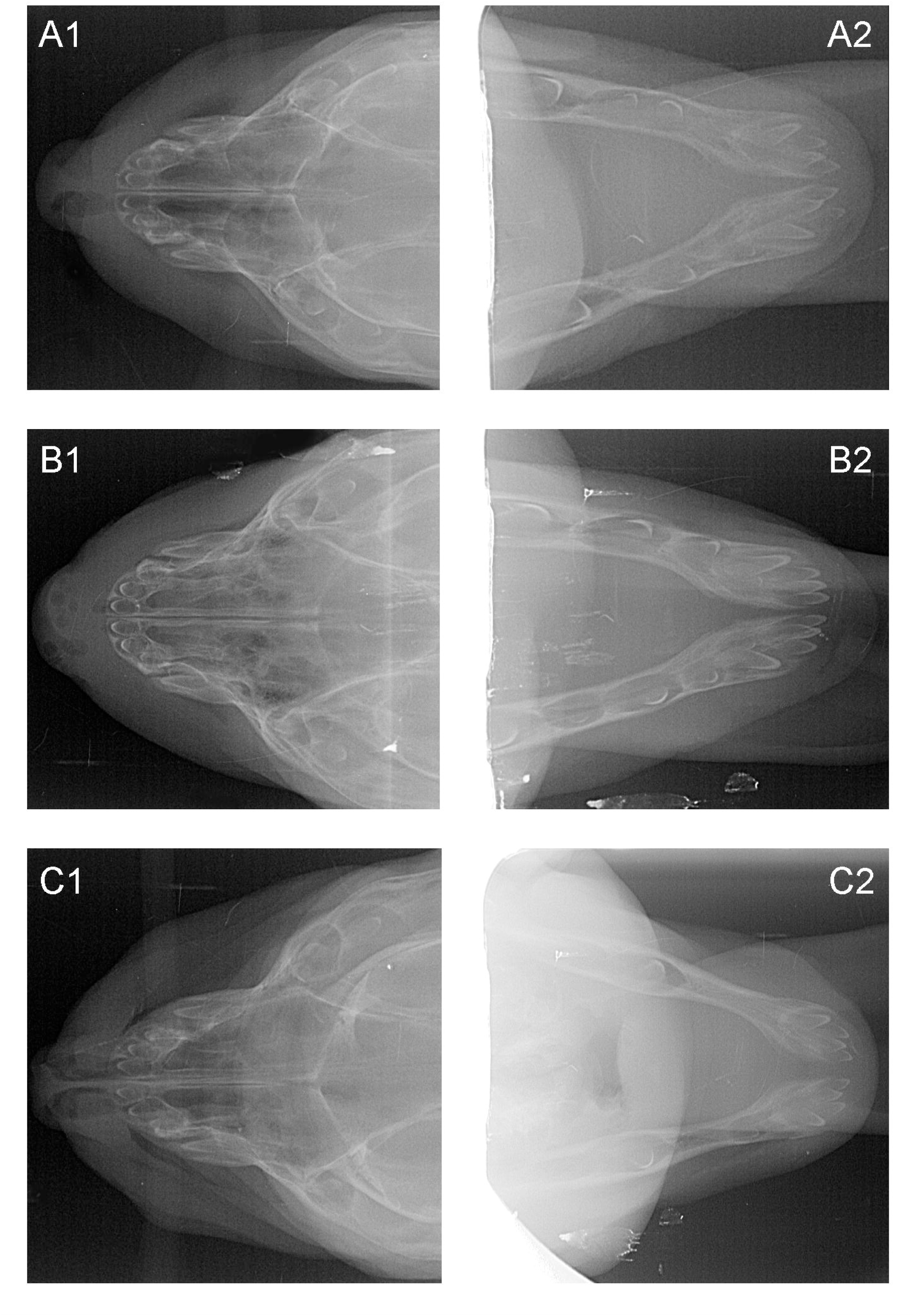
Dental imaging for dogs B, C and D. Depicted are the mandibular (right) and maxillar (left) X-rays of three of the affected male puppies. A healthy puppy should have the following developing teeth in the upper and lower jaw at the age of eight days: six incisors, two canines, four premolars and two molars. In the upper jaw of dog B (A1), two premolars are missing, the lower jaw (A2) has a full bite but with irregularly spaced teeth. Dog C has a full bite in both jaws and additional premolar in the lower jaw (B2). Dog D has two missing premolars in the upper jaw (C1) and three missing premolars in the lower jaw (C2).

### Pedigree analysis

The pedigree of the litter with the affected puppies was traced back for six generations and phenotype data were supplemented for these dogs. The dam and sire of the affected puppies were unaffected. A litter of maternal half-siblings was inconspicuous as well as all littermates of the dam and granddam and further siblings of all progeny of ancestors. The maternal granddam (dog F) and maternal great-granddam (dog G) had healthy litters from different sires. An autosomal recessive or dominant inheritance was ruled out due to normal parents and ancestors as well as no obvious inbreeding in the six generations. The mode of inheritance suggests an, X-recessive transmission through the dam or the germline of either grandparent (Figure 2).

### Whole genome sequencing, Sanger sequencing and mutation analysis

Whole genome sequencing of dog A revealed a 1-bp deletion in exon 6 within the *EDA* gene (CM000039.3:g.54509504delT, c.458delT). Variants in other ED candidate genes (Table S1) did not fulfill the search criteria. Poly-Phen2 predicted this mutation to be probably damaging with a score of 1.000 (sensitivity: 0.00; specificity: 1.00) using the HumDiv model. Sanger sequencing of DNA-samples of the affected puppy, dam and granddam (dogs A, E and F) confirmed dog A as hemizygous, his dam (dog E) as heterozygous and the granddam (dog F) as wildtype (Figure 5). A cDNA Sanger sequence of dog E was hemizygous wildtype. This *EDA*-1-bp deletion was predicted to lead downstream to 21 altered amino acids (aa) and a premature stop codon in the canine *EDA* transcripts EDA-201 and EDA-202 (Figure 6). The protein sequence is predicted to lack 85 aa and thus be shortened from 258 to 173 aa (p.H153fsX173). According to the canine reference genome assembly CanFam3.1, the frameshift mutation is localized at amino acid position 153 and therefore in the tumor necrosis factor-like domain or tumor necrosis factor domain of ectodysplasin.

**Figure 5:**
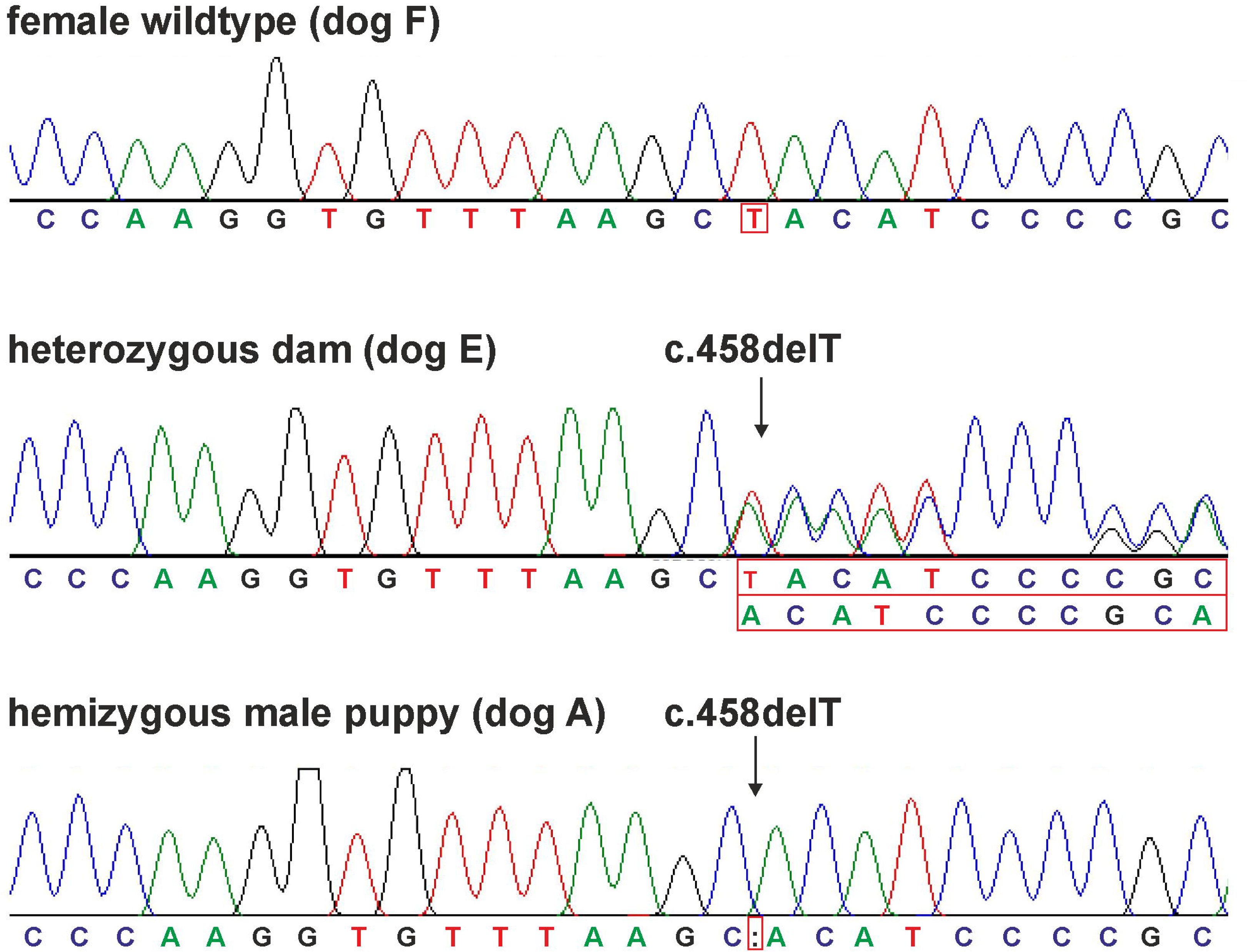
Sanger sequencing chromatograms of dogs A, E and F. The granddam (dog F) was homozygous wildtype. At position *EDA*:c.458 the wildtype allele T is present. One of the affected male puppies shows at position *EDA*:c.458 deletion of the allele T (dog A). His dam (dog E) is heterozygous resulting in a shift of the sequence with the PRJEB27789 variant.

**Figure 6:**
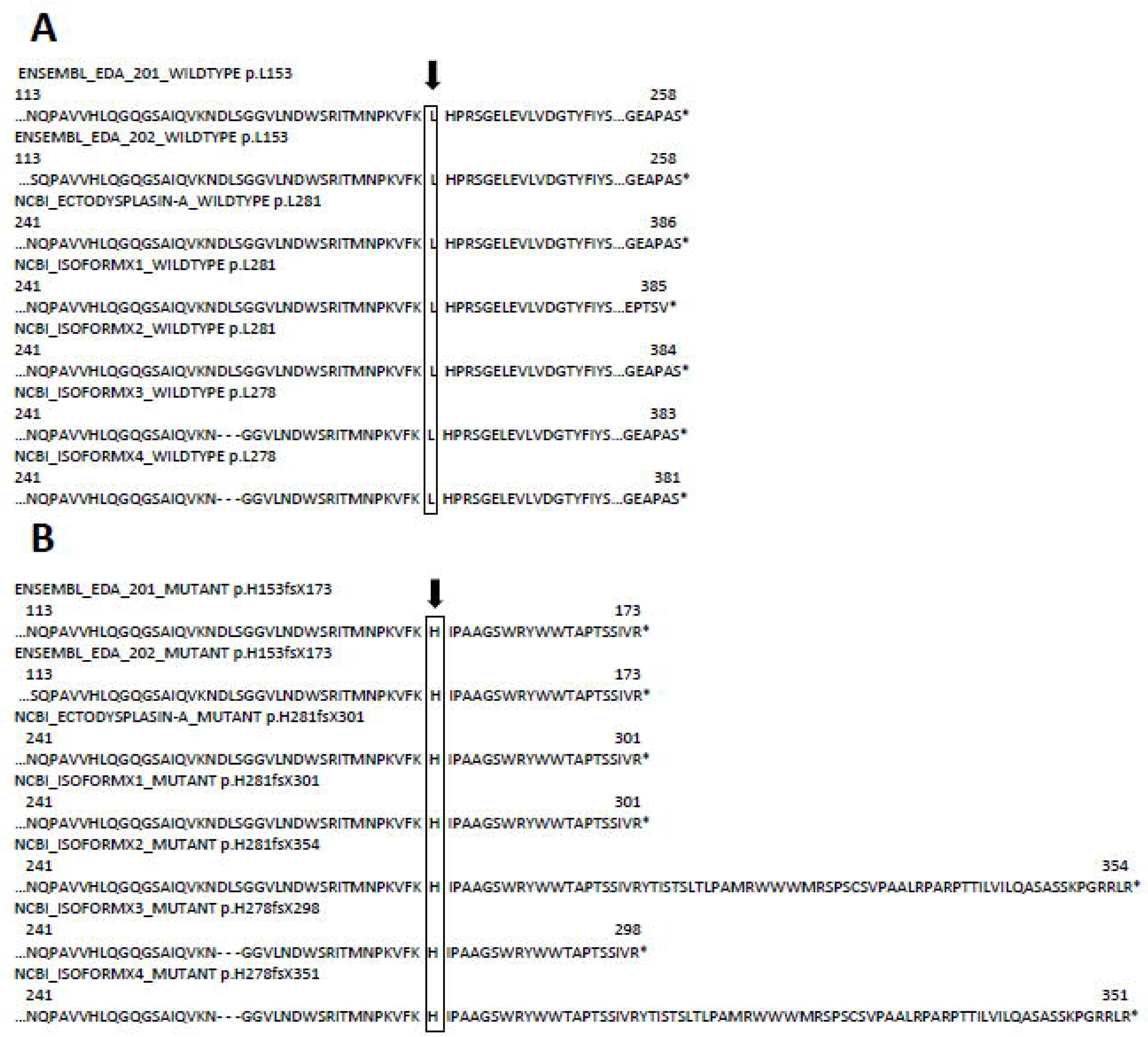
Given are the known ectodysplasin-A-protein isoforms of the canine *EDA* gene and the predicted effect of the *EDA*:c.458delT frameshift mutation on their proteins. First, we compared the various wildtype protein sequences as stated by the databases Ensembl and NCBI (A). Then we present the respective predicted mutant protein sequences (B). The frameshift mutation was predicted to lead to 21 altered amino acids and a premature stop codon in the *EDA* transcripts EDA-201 and EDA-202 (Ensembl). The produced proteins therefore would lack 85 amino acids. The database NCBI (https://www.ncbi.nlm.nih.gov/) lists another five ectodysplasin-A isoforms, which would also be shortened by 85 to 30 amino acids. The stated protein sequences of both databases differ. EDA-201 and EDA-202 have the same length of 258 amino acids, but differ on 10 amino acids. Ectodysplasin-A as stated by NCBI has a sequence identical to EDA-201 but is 128 amino acids longer, as the predicted transcription starts earlier. The other isoforms of ectodysplasin-A are one to five amino acids shorter and with a few non-matching amino acids. According to the canine reference genome assembly CanFam3.1, the frameshift mutation is localized at amino acid position 153 (281 resp. 278 for the NCBI-isoforms) and therefore in the tumor necrosis factor-like domain or tumor necrosis factor domain of ectodysplasin. In each case, the mutation leads to an altered and shortened protein sequence, thus rendering the tumor-necrosis-factor-domain of the ectodysplasin-A isoforms useless.

### Validation

PCR-RFLP revealed all four affected littermates as hemizygous mutant. The dam (dog E) was heterozygous and the maternal granddam (dog F) and maternal great-granddam (dog G) were wildtype. Female progeny of the grandsire were wildtype. The mildly hypotrichotic female siblings of the cases (dogs I and J) were heterozygous, and the female littermate with a full coat (dog H) was wildtype. A summary of all available genotypes of the family is given in Table 1. Furthermore, the *EDA*-variant was homozygous wildtype in 37 unrelated shorthaired standard Dachshunds as well as in other Dachshund breeds including shorthaired miniature (n=1), longhaired rabbit (n=6), longhaired standard (n=4), longhaired miniature (n=14), wirehaired rabbit (n=9), wirehaired standard (n=59) and wirehaired miniature (n=35) as well as in the across-breed screen for 34 different dog breeds (Table S3). We found a perfect association of the CM000039.3:g.54509504delT variant with the affected phenotype and confirmed X-linked inheritance.

**Table 1.** Genotyping results for the CM000039.3:g.54509504delT variant of the affected Dachshund family and reference dogs. The Dachshund breed, the number of individuals per breed, and the respective sex and genotype are stated. For the affected dachshund family, the predicted genotype is also stated, when it could be deducted from phenotype and/or mode of inheritance.

## Discussion

In this study, we provide an in-depth description of an XLHED case in a litter of shorthaired standard Dachshunds. The diagnosis XLHED was based on the specific phenotype of the puppies, their pedigree and a mutation analysis using WGS-data. The most obvious signs for XLHED were the partial alopecia with a consistent pattern among all four male cases and the tooth anomalies. This alopecia pattern is identical to those of dogs with confirmed cases of XLHED (Casal et al. 1997; Waluk et al. 2016) and also consistent with many other cases where XLHED is suspected (Moura and Cirio 2004), namely a bald forehead, abdomen and dorsal pelvic area. Secondly, the observed tooth anomalies are typical findings in XLHED (Lewis et al. 2010). Whether the remaining teeth were misshapen could not be determined on the dental X-ray of the eight-day old pups. Subsequently, the evaluation of the permanent dentition is also not applicable. The fact that one of the puppies (dog C) does not exhibit detectable tooth anomalies is not contradictory to the diagnosis of XLHED. In one case, out of 17 examined XLHED affected dogs, one had no missing teeth (Lewis et al. 2010), and in another it was stated that there were no tooth abnormalities in the two XLHED-affected, 15 month old miniature poodles (Selmanowitz et al. 1970a), suggesting some variance in the severity of affectedness. Another organ system affected in ED are exocrine glands (Cluzeau et al. 2011), which is given in our case. The examined puppy was missing eccrine sweat glands, tracheal and esophageal glands. The absent tracheal glands have led to decreased mucociliary clearance and upper respiratory tract infections, which presented as the nasal discharge and respiratory distress the male puppies were affected with. Though they did not suffer from bronchopneumonia, the signs were severe enough to prompt euthanasia of the affected male puppies.

Pedigree analysis revealed that the mode of inheritance is recessive, since the litter was born from unaffected parents. A X-chromosomal inheritance was assumed due to only male affected offspring. Thus, the most likely mode of inheritance is X-recessive with the dam as a heterozygous carrier of the mutation as confirmed through the PRJEB27789 variant. The maternal granddam was homozygous wildtype. A DNA sample could not be obtained from the grandsire, but his normal phenotype is consistent with a X-chromosomal inheritance. We also did not identify any other offspring of granddam and -sire as carriers of the *EDA*-mutation. That source of the *EDA*-mutation may be most likely in the germline of the granddam, or in an early embryonic stage of the dam (dog E). A germline mutation in the grandsire seems less likely, as first and later offspring generations were normal and ED phenotypes were not reported from this grandsire line. In case of a germline mutation of the granddam, all somatic and germline cells of the dam harbor the mutation. We cannot exclude that the dam is a mosaic of normal and mutated cells, supporting the hypothesis of an early embryonic mutation before germ cells differentiated.

With the dam being an X-heterozygous, asymptomatic carrier one would expect that 50% of her male descendants would be affected and 50% of her female offspring carriers. The actual ratio of healthy versus affected male descendants of the dam is 1:4, which is still within statistical limits given the small sample size. Furthermore two of her six daughters (dogs I and J) were identified as carriers. The two carrier females exhibited sparse hair on the frontotemporal area, ears and ventral chest. The dentition of the dam and her carrier daughters at the age of eight weeks was inconspicuous. It is described that carrier females often suffer mild symptoms, too (Kerr et al. 1966; Söderholm and Kaitila 1985; Clarke et al. 1987; Casal et al. 1997). The variance in the severity of signs may be attributed the random inactivation of one X-chromosome in female mammals (lyonization) (Moura and Cirio 2004; Lexner et al. 2008; Vogel and Motulsky 2013). Alternatively, if the dam was a mosaic and the amount of cells carrying the mutation would be low (also called a low-level mosaicism), this would lead to the same phenotype as a skewed X-inactivation to the healthy side. Low-level mosaicism in the dog is a rare condition, but may be underdiagnosed (Switonski et al. 2003).

For the mutation detection in the WGS data, the filtering criteria left only *EDA* as a candidate gene. This was expected, since the other most common ED candidate genes do not match our observed phenotype and mode of inheritance. *EDAR* and *EDARADD* mutations would cause a clinically indistinguishable phenotype, but are inherited as autosomal, and *WNT10A* also causes a distinct phenotype with nail dystrophy (Cluzeau et al. 2011). Mutations in further ED candidate genes *TNF receptor associated factor 6* (*TRAF6), Inhibitor Of Nuclear Factor Kappa B Kinase Subunit Gamma (IKBKG,* formerly *NEMO), NF*κ*B inhibitor* α *(NFKBIA*) and *NF*κ*Bs* also cause immunodeficiency (Cui and Schlessinger 2006) and variants with effects on protein function were not identified in these candidates. Subsequently the CM000039.3:g.54509504delT variant was the only one that met all conditions and was validated.

The mutation c.458delT causes a frameshift, leading to altered aa and a premature stop codon and thus shortening of all canine ectodysplasin-A isoforms. The mutation was predicted to affect the tumor-necrosis-factor domain of the protein, thus probably rendering this domain useless (Schneider et al. 2001). Depiction of the mutated mRNA by Sanger sequencing a cDNA-PCR product of the heterozygous dog E was attempted, but apparently, only the mRNA from the intact chromosome was sequenced, so the Sanger sequence came out as (hemizygous) wildtype. We therefore speculate that in case of a mosaicism, it is possible that the hair collection site (the tail) consisted only of normal cells. Another possibility is that the mutant mRNA is produced, but quickly detected as faulty by the organism and therefore degraded. It is possible that the small fragments of the degraded mRNA showed up as mild interference in the chromatogram of the Sanger sequence. The degradation of mRNA with premature stop codons is a well-known phenomenon, called nonsense-mediated decay (Wagner and Lykke-Andersen 2002). The degradation can be initiated by the removal of the mRNA poly-A tail, which provides access for degradative enzymes to the body of the mRNA (Wilusz et al. 2001). If the dam’s X-inactivation was skewed to the healthy side, so that more of the mutant than of the healthy chromosome was inactivated, this would leave a large enough amount of functional ectodysplasin for her to not develop ED signs. Alternatively, if the mutant *EDA* was transcribed, the protein would be nonfunctional because of the damaged TNF-domain.

The TNF-domain is crucial to the function of ectodysplasin-A (Schneider et al. 2001). The TNF-domain of this type II membrane protein is located extracellular (Ezer et al. 1999). From there the TNF-domain is cleaved to produce a soluble ligand that forms trimers and interacts with its respective receptors (Elomaa et al. 2001; Chen et al. 2001). One isoform of ectodysplasin-A, EDA-A2, is specific to the receptor EDA2R (Yan et al. 2000). This pathway can activate NF-κB in vitro (Yan et al. 2000; Sinha et al. 2002), but not in-vivo (Gaide and Schneider 2003; Mustonen et al. 2003). Furthermore, the role of EDA2R signaling seems expendable in skin development, as *EDA2R* knockout mice do not show an ED phenotype (Newton et al. 2004). EDA-A1, however, is well known for binding to its receptor EDAR (Yan et al. 2000). EDAR in turn binds to its cytoplasmic signaling adaptor EDARADD (Headon et al. 2001), which via TRAF6 activates the IκB kinase (IKK) complex (Morlon et al. 2005). From there on, apparently the IKBKG/IkBα-dependent NF-κB cascade is initiated (Cui and Schlessinger 2006; Weih and Caamaño 2003). The NF-κB pathway is crucial for the development of skin appendages. Thus it is plausible, that a mutation in the TNF domain of ectodysplasin-A, leads to impaired binding of both isoforms to their receptors (Schneider et al. 2001), subsequently leading to a disrupted EDA/NF-κB signaling pathway and manifesting itself in XLHED. In conclusion, this study demonstrates the first XLHED cases in the shorthaired standard Dachshund, most likely due to an *EDA* germline mutation in the dam or maternal granddam. The PRJEB27789 variant is predicted to cause a nonfunctional protein due to a damage of the TNF-domain needed for the *EDA* signaling pathway. Variance in affectedness of carrier females was obvious and possibly attributable to a skewed X-inactivation, or, in case of the dam, to low-level mosaicism.

## Acknowledgements

The authors would like to thank the owners of the dogs included in the study for donating samples and sharing information about their dogs, particularly the Dachshund breeder for reporting the case and providing us with pedigree data. Special thanks go to J. Wrede, M. Drabert, H. Klippert-Hasberg and N. Wagner for their expert technical assistance. We gratefully acknowledge support from the North-German Supercomputing Alliance (HLRN, Hanover) for HPC-resources that contributed to the research results.

## Supplementary Files

**Table S1** Candidate gene list analysis. Genes potentially involved in ectodermal dysplasia were identified using NCBI Gene database. The canine chromosome position, genes, gene ID and name are shown.

**Table S2** Primer sequences used for validation of the mutation detected by whole genome and Sanger sequencing. The variant CM000039.3:g.54509504delT, c.458delT was validated by the use of restriction fragment length polymorphism (RFLP). Primer pairs, amplicon size (AS) in base pairs (bp), annealing temperature (AT), restriction enzyme and incubation temperature (IT) are given. Also depicted are the three primers used for cDNA Sanger sequencing.

**Table S3** Genotyping results for the CM000039.3:g.54509504delT variant of reference dogs of other breeds. The dog breed, the number of individuals per breed, and the respective sex and genotype are stated. 70 dogs of 34 breeds were tested in total.

